# Tonotopic mapping in children shows an overrepresentation of high frequencies in autism

**DOI:** 10.64898/2026.10.06.757151

**Authors:** Bosi Chen, Alexandra Boxberger, Judy Mahmalji, Tess Do, Meagan Herrera, Sabrena Tuy, Giovanna Arantes De Oliveira Campos, Martin I. Sereno, Inna Fishman, Annika C. Linke

## Abstract

Altered sensitivity to sounds despite normal peripheral hearing is common in neurodevelopmental disorders but the underlying mechanisms remain poorly understood. Here we investigate tonotopic organization in 86 children (6-11 years) with autism spectrum and related neurodevelopmental disorders as well as typically developing (TD) peers, using a child-friendly, passive listening, functional magnetic resonance imaging design. In TD children, tonotopic maps in primary auditory cortex resembled the typical high-low-high frequency gradients previously reported in adults, while children with autism showed an overrepresentation of high frequencies consistent with what has been observed in animal models of autism. Higher frequency tuning further correlated with greater caregiver-reported sound but not touch sensitivities. Importantly, tonotopy is a core organizational feature of auditory cortex that develops early in life and remains relatively stable after the critical period of heightened plasticity ends. Thus, these results suggest that atypical auditory processing in autism may result from early disruption to tonotopic map formation. Longitudinal neuroimaging studies are needed to understand the genetic and environmental causes of altered auditory cortex topology and its long-term consequences for higher-level auditory, language, and social communication development.

## Introduction

Topological maps reflecting orderly spatial representations are one of the core features of brain organization. While sensory maps, like retinotopy in visual cortex and tonotopy in auditory cortex, have been characterized extensively using functional magnetic resonance imaging (fMRI) in neurotypical adults^1–3^, they have rarely – and, in the case of tonotopy, not at all – been studied in children. This gap is notable given that the auditory system is one of the first to mature, making it particularly vulnerable to disruption early in life with potential long-term consequences for language and social development.

Understanding the neural mechanisms underlying atypical sound processing in neurodevelopmental disorders is crucial given that it affects up to 90% of children with autism spectrum disorders (ASD)^4^ and ∼50% of children with attention deficit hyperactivity disorder (ADHD)^5^. Increasing evidence points toward a central disruption of auditory processing underlying altered sensitivity to sounds in these neurodevelopmental disorders^6,7^. For example, in one study of school-age children with ASD who presented with perceived hearing difficulties, almost 90% showed deficits on tests of speech recognition in noise, auditory filtering, or binaural integration and attention, despite normal hearing thresholds^8^. Similarly, neuroimaging studies have consistently shown atypical cortical responses to sound in individuals with autism and normal peripheral hearing^9–12^. In humans, hearing begins in utero, with the brainstem and auditory cortex achieving maturity necessary for processing sounds around 28 weeks of gestation^13^. Rodent models show that tonotopic maps form around the time of hearing onset, according to a genetically determined process, and undergo rapid refinement during a short critical period after which maps remain mostly stable throughout life^14,15^. Genetic predisposition and known environmental risk factors for ASD, such as complications during gestation or birth^16,17^, can interfere with this development^18^. In a valproic acid (VPA) mouse model of autism, tonotopic maps show distortions with an overrepresentation of high frequency tones^19^. Disrupting the development of primary auditory cortex has long-term consequences in animal models, such as resulting in altered vocalization^20^. As such, it is crucial to investigate how common are alterations of auditory topological maps in autism and related neurodevelopmental disorders, and how this relates to heightened sensitivity to sounds experienced by children with these conditions. The current study is the first to show that tonotopic mapping is possible in children using a short and engaging fMRI protocol and the first to compare tonotopic organization in children with ASD and related neurodevelopmental disorders (NDD) to age- and sex-matched typically developing (TD) children.

## Materials and Methods

One hundred children between 6 and 11 years of age (n_TD_=41, n_ASD_=33, n_NDD_=26) participated in ongoing fMRI studies (conducted between 2021 and 2026) investigating neural mechanisms of atypical auditory processing at San Diego State University. The research protocol was approved by the institutional review board of San Diego State University and parental consent and child assent were obtained prior to participation. Upon enrollment, all children with (or suspected to have) ASD, ADHD, or another developmental condition, underwent full diagnostic evaluation, using standardized measures (e.g., Autism Diagnostic Observation Schedule, Second Edition [ADOS-2], Conners-3) in combination with expert clinical judgment, based on the DSM-5 criteria. Caregiver and teacher forms (including the Child Behavior Checklist, Sensory Processing Measure) were also administered. For inclusion in the TD group, children were required to have below clinical cutoff scores on the ASD screener (Social Communication Questionnaire [SCQ]) and on Conners-3, and to demonstrate overall cognitive abilities falling no more than 1.5 standard deviation (SD) below the normative mean for their age. Exclusion criteria are summarized in the Supplementary Materials. All children were reported to have normal peripheral hearing.

For tonotopic mapping, two 5-minute fMRI runs were acquired on a 3T Siemens Prisma scanner at the SDSU MRI Center using a multiband EPI sequence (acceleration factor=4, TR=1125ms, TE=33ms, flip angle=65°, voxel size=2.5mm isotropic, 56 slices, 264 volumes including 8 dummy scans discarded during pre-processing). Upward and downward sweeping pure tone sequences (low-frequency extremes: 130-175 Hz; high frequency extremes: 3000-4000 Hz) were created with pure tones randomly sampled within 1/2-octave intervals and presented at a rate of 1 octave every 8.72s. Each fMRI run included six upward or downward sweep sequences (48s long each) with no gap between successive sweeps delivered via active noise-cancelling headphones (OptoActive II, >30dB noise reduction). A silent, child-friendly animated movie without any verbal interactions between characters was played in the background to increase engagement and reduce head motion. MATLAB with Psychtoolbox was used to synchronize EPI acquisition, auditory stimuli, and movie onset. High-resolution T1-weighted anatomical scans (1mm^3^ voxel size) were also collected for each child.

MRI data were preprocessed using SPM12 and the CONN toolbox (v21), including rigid-body realignment, normalization to the Montreal Neurological Institute template, high-pass filtering (0.008 Hz), and nuisance regression to remove physiological and motion confounds. Images were spatially smoothed with a 6mm full-width at half-maximum (FWHM) Gaussian kernel and voxel time series were converted to percent signal change.

Following methods described by Langers et al.^21^, linear regression models were used to fit a cosine and sine function modelling the sinusoidal responses to the pure tone sweeps. The sign of the sine function was flipped for the runs with downward sweeps. Regression coefficients were averaged between the two runs and converted to polar coordinates, reflecting response amplitude 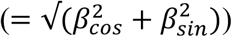 and frequency tuning (phase = arctan(*β*_*sin*_ /*β*^*cos*^)). The median amplitude and frequency tuning were then calculated for voxels in regions A1, R, and RT combined (i.e., primary auditory cortex, as defined by Sereno et al.^22^) for each hemisphere. General linear models were used to test differences in frequency tuning by diagnostic group, age, biological sex (male, female), and group-by-sex interactions, separately for left and right primary auditory cortex, with mean head motion during the scan as a covariate.

The Sensory Processing Measure (SPM) Hearing T-score was used in general linear models to assess relationships between average frequency tuning in the primary auditory cortex (per hemisphere) and auditory processing symptoms. The SPM, Home Form is a standardized caregiver questionnaire that yields normalized T-scores (mean=50, SD=10) of sensory processing across different sensory systems as well as a Total T-score. A T-score ≥ 60 indicates sensory processing difficulties. Due to floor effects observed in TD children (Hearing T-score: mean=50.4, SD=9.4 in the current cohort), analyses were only restricted to children with ASD (SPM available for n_ASD_=21/28, 11 female; mean=64.1, SD=8.1) and those with other NDD (SPM available for n_NDD_=20/20, 10 female; mean=56.4, SD=10.5). Mean head motion during scan and sex were included as covariates since, within the combined ASD and NDD group of children, Hearing T-scores differed significantly (*t*(39)=2.2, *p*=.03) between males (mean=58.5, SD=10.5) and females (mean=65.8, SD=9.9), see Figure S1.

## Results

Data from 14 children (n_TD_=3, n_ASD_=5, n_NDD_=6) were excluded due to excessive head motion during the scan (n=2) or not completing the MRI scans (n=12, including two with MRI contraindications). Consequently, high-quality tonotopic mapping fMRI data were available for 38 TD children (20 female), 28 children with ASD (11 female), and 20 children with diagnoses of other neurodevelopmental disorders (NDD; 10 female, including ADHD (n=14), oppositional defiant disorder (ODD; n=2), intellectual disability (n=1), or siblings of a child with ASD (n=3)). The three groups did not differ significantly in age, sex, gestational age at birth, or head motion during fMRI scans (see Table 1). Response amplitude in left and right primary auditory cortex did not differ significantly between children with ASD and those with other neurodevelopmental disorders (NDD) or TD children, though left primary auditory cortex showed slightly lower amplitude in the NDD group compared to TD children (*t*(81)=2.4, *p*=.049). These results suggest that any potential group differences in frequency tuning are unlikely attributable to differences in demographic characteristics, fMRI data quality, or overall activation magnitude in response to the passive listening paradigm.

**Table 1.** Demographics for children with successful fMRI scans by diagnostic group.

|  | TD<br>(n=38) | ASD<br>(n=28) | NDD<br>(n=20) | Group comparison |
| --- | --- | --- | --- | --- |
| Sex<br>(% female) | 53% | 39% | 50% | $\chi^2 = 1.2, p=.55$ |
| GA at birth (weeks) | 39.6 ± 1.1<br>[37-42] | 39.7 ± 1.7<br>[34-43] | 38.7 ± 2.7<br>[29-41] | $F(2,75)=1.9, p=.15$ |
| Age at MRI scan<br>(years) | 8.2 ± 1.6<br>[6-11.9] | 8.5 ± 1.9<br>[6-11.7] | 8.5 ± 1.7<br>[6.3-11.6] | $F(2,83)=0.2, p=.8$ |
| Ethnicity<br>(% Hispanic or Latino) | 28% | 52% | 37% | $\chi^2 = 3.7, p=.16$ |
| Household Income* | 8.4 ± 2.0<br>[2-10] | 7.6 ± 1.9<br>[5-10] | 8.0 ± 2.1<br>[3-10] | $F(2,67)=1.2, p=.31$ |
| Maternal Education<br>Level** | 5.6 ± 1.1<br>[3-7] | 4.0 ± 2.0<br>[1-7] | 4.8 ± 1.8<br>[1-7] | $F(2,71)=6.2, p=.003$ |
| ADOS-2<br>Comparison<br>Score *** | n/a | 6.5 ± 2.0<br>[1-10] | 3.6 ± 2.4<br>[1-7] | $t(39)=4.4, p<.001$ |
| ADHD<br>Diagnosis (n) **** | n/a | 11 | 15 | -- |
| FSIQ ***** | 111 ± 13.2<br>[81-137] | 103 ± 18.4<br>[71-130] | 105 ± 16.7<br>[65-131] | $F(2,80)=2.0, p=.14$ |
| SPM Hearing<br>T score | 49.7 ± 8.7<br>[43-70] | 65.9 ± 11.2<br>[43-80] | 57.0 ± 9.4<br>[43-71] | $F(2,64)=16.1, p<.001$ |
| Mean Head Motion<br>(fMRI) | .23 ± .08<br>[.14-.47] | .25 ± .07<br>[.12-.44] | .22 ± .07<br>[.15-.41] | $F(2,83)=1.6, p=.21$ |

\* Household income was coded on a 1-10 scale (ranging from 1= <\$10,000 to 10= >\$200,000). Information about household income was missing for n=8 TD children and n=8 children with ASD.
\*\* Maternal Education Level was coded on a 1-7 scale (ranging from 1=High School/GED or equivalent to 7=Professional Degree). Information about maternal education level was missing for n=5 TD children and n=7 children with ASD. Maternal education level was significantly lower in children with ASD compared to TD children ( $p_{\text{tukey}}=.004$ ) but not significantly different compared to children with other neurodevelopmental disorders ( $p_{\text{tukey}}=.23$ ).
\*\*\* The ADOS-2 was only administered for 12/20 children in the NDD group.
\*\*\*\* In the ASD group, this reflects co-occurring ADHD diagnosis.
\*\*\*\*\* The Differential Ability Scales (DAS, n=10) or WASI-II were administered to estimate Full Scale IQ (DAS General Conceptual Ability, FSIQ-4 Composite scores). FSIQ scores were missing for n=2 TD children and n=1 child with ASD.

Significant main effects of diagnostic group were observed for frequency tuning in both left (*F*(2,78)=5.3, *p*=.007) and right (*F*(2,78)=3.8, *p*=.027) primary auditory cortex (Figure 1A, Figure S2). Post-hoc tests showed that these effects were driven by higher frequency tuning in the ASD compared to the TD group (see Figure 1B). There were no significant main effects of sex or sex-by-diagnosis interactions, nor were there significant associations with age or head motion. Additional post-hoc analyses showed no significant effects of maternal education level or gestational age at birth (see Supplementary Material).

**Figure 1.**
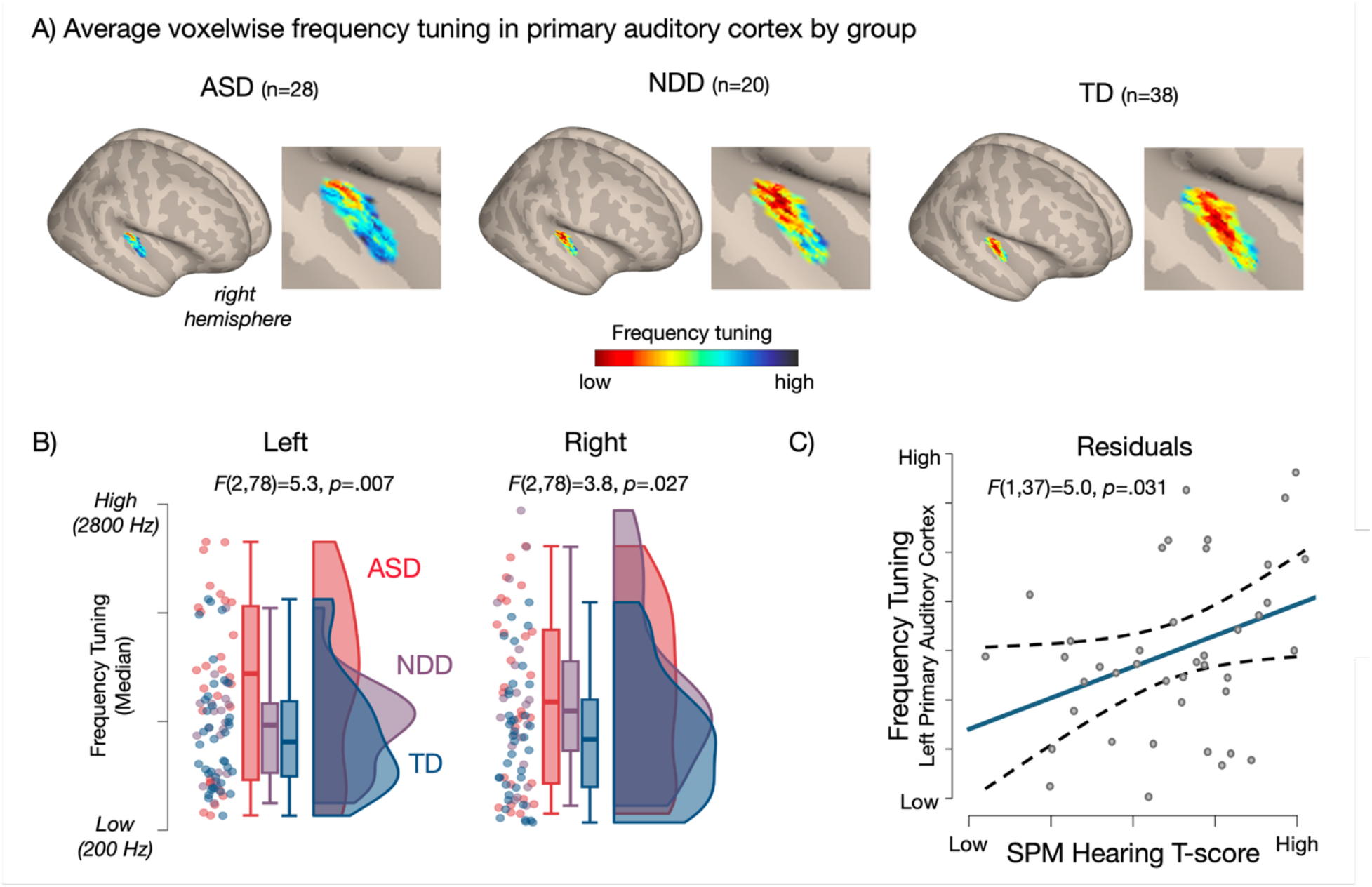
**A)** Maps of average frequency tuning for each voxel in right primary auditory cortex for each diagnostic group. Hot (red and yellow) colors reflect tuning to lower and cold (blue) colors to higher frequencies. An overrepresentation of voxels tuned to high frequencies is evident in the ASD group. Maps in the NDD and TD groups resemble the typical high-low-high gradient previously reported in adults. Maps for left primary auditory cortex are shown in supplementary Figure S2. **B)** Frequency tuning in left and right primary auditory cortex shows a significant main effect of diagnostic group. Pairwise post-hoc tests indicate that this is driven by significantly higher frequency tuning in the ASD compared to TD group (left: *p*_tukey_=.008, right: *p*_tukey_=.03). The pairwise post-hoc test was also significant for left primary auditory cortex when comparing the ASD to NDD group (*p*_tukey_=.04). Pairwise comparisons between the NDD and TD group were not significant. **C)** Elevated auditory processing difficulties as measured by the Sensory Processing Measure (SPM, Home Form) Hearing subscale (T-scores) are associated with higher average frequency tuning in left primary auditory cortex in children with ASD and with other NDD. Model controlling for sex and head motion during the fMRI scan; residual plot (with unitless axes) is shown. SPM Hearing T-scores ranged from 43 to 80 (mean=61.5, SD=10.8) with T-scores above 60 indicating sensory processing difficulties (T=60-69: “some problems”, and T>70: “definite dysfunction”; n=24/41 children with ASD or with other NDDs had T>60).

Among children with ASD or other NDD, higher frequency tuning in the left primary auditory cortex was associated with elevated SPM Hearing subscale T-scores (*F*(1,37)=5.0, *p*=.03, Figure 1C). There was no significant relationship between auditory sensory symptoms and right primary auditory cortex frequency tuning (*F*(1,37)=2.0, *p*=.16). To test the specificity of this association to auditory processing, analyses were repeated with the SPM Touch T-scores, which revealed no significant effects (left: *F*(1,37)=.92, *p*=.34, right: *F*(1,37)=.71, *p*=.41).

## Discussion

This study demonstrates that tonotopic mapping using a short and engaging fMRI paradigm that requires only passive listening is feasible in children, including those with neurodevelopmental disorders. While tonotopy has not been previously characterized in younger children, the absence of an age effect in our sample (ages 6-11 years) aligns with critical periods for auditory cortex development established from animal and human studies. Research in animal models has shown that tonotopic organization in primary auditory cortex is largely established during early postnatal development and stabilizes after the closure of the critical period^23–25^. Human studies similarly demonstrate that auditory cortex myelination and synaptogenesis reach adult-like levels by early childhood^26^. Specifically, studies of children receiving cochlear implants have demonstrated that the central auditory system is maximally plastic during the first 3.5 years of life, with plasticity declining substantially by age 7 years^27^. After the critical period closes, tonotopic organization typically only undergoes changes in extreme circumstances (e.g., prolonged and intense exposure to the same frequency stimulus^28^) or in advanced age (e.g., in response to sensorineural hearing loss^29^).

In TD children, tonotopic maps in primary auditory cortex resembled the canonical high-low-high gradients reported in adults in previous studies, with average frequency tuning being significantly lower than in children with ASD. The relative overrepresentation of high frequencies observed in children with ASD aligns with findings from a VPA mouse model of autism^19^. In rodents, prenatal VPA exposure also results in atypical vocalizations^20^ (in neonatal, adolescent and even adult rat offspring), suggesting that altered tonotopic organization may have functional implications for auditory processing and communication, across the animal lifespan. The preferential tuning toward higher frequencies in humans with ASD could reflect an expanded cortical area devoted to high-frequency processing, which may contribute to the auditory hypersensitivities commonly reported in ASD. In the current study, higher frequency tuning in the left auditory cortex was associated with greater auditory processing difficulties as assessed with the Sensory Processing Measure (SPM) in children with ASD and other NDD. This finding provides empirical support for a link between atypical tonotopic organization and real-world auditory processing challenges. Notably, this relationship was specific to auditory processing difficulties, suggesting that the observed tonotopic alterations have domain-specific functional consequences. Future neuroimaging studies in longitudinal cohorts will be needed to understand the genetic and environmental factors contributing to altered auditory cortex topology and its long-term consequences for higher-level auditory, language, and social communication development.

Importantly, although our results suggest that an overrepresentation of high frequencies in auditory cortex is more prevalent in children with ASD, this does not imply that it is specific to ASD, particularly given the relatively small sample size of children with other neurodevelopment disorders (predominately ADHD) included in the current analyses. Atypical tonotopic organization has also been reported in other clinical populations, including, notably, in adults with schizophrenia who experience recurrent auditory hallucinations^30^. Future studies with larger and more diverse cohorts as well as dimensional approaches to symptom characterization will help identify clinical and behavioral factors most strongly associated with altered tonotopic map development and clarify how these early sensory differences interact with other risk factors for neurodevelopmental and neuropsychiatric disorders.

## Data Availability

Data used for the analyses presented in this manuscript are available to researchers upon request for those participants who provided data sharing consent.

## Acknowledgements

We thank Michaela Cordova, PhD (San Diego State University) and Sara Bock (San Diego State University) for assistance with data collection, training of students and staff and contributions to the project implementation. Our strongest gratitude goes to the children and families who so generously dedicated their time and effort to this research.

