## Supplementary Material for "Tonotopic mapping in children shows an overrepresentation of high frequencies in autism"

### Exclusion criteria

Exclusion criteria for study participation included the following: reported neurological disorder (other than ASD, e.g., epilepsy), syndromic forms of ASD (e.g. fragile X or Rett syndrome), any clinically significant metabolic abnormality, central nervous system (CNS) infection or gross CNS injury (e.g., cerebral palsy or brain injury), or any fMRI contraindications. For the TD group, further exclusion criteria included prematurity (<36 weeks of gestation), first-degree relative with ASD, intellectual disability, other heritable psychiatric or neurological disorders, scores on the ASD screener (Social Communication Questionnaire [SCQ]) above clinical cutoff, and overall assessed cognitive abilities lower than 1.5 SD below the normative mean for their age. Co-occurring conditions (e.g., ADHD) in children with ASD were not exclusionary (that is, children with autism and co-occurring ADHD or ODD were included in the ASD group). On the other hand, no children in the NDD group had ASD.

### Effects of Maternal Education Level and Gestational Age at Birth

The main effect of group was also significant when including maternal education level (left primary auditory cortex: $F(2,67)=5.1$, $p=.009$, right: $F(2,67)=3.1$, $p=.05$), or gestational age at birth (left primary auditory cortex: $F(2,71)=5.3$, $p=.007$, right: $F(2,71)=4.1$, $p=.02$) as covariates (models testing for a main effect of group, while controlling for age, sex, and mean head motion).

The main effects of maternal education level (left: $F(1,67)=0.2$, $p=.62$, right: $F(1,67)=0.3$, $p=.61$) and gestational age at birth (left: $F(1,71)=.5$, $p=.5$, right: $F(1,71)=3.7$, $p=.06$) on frequency tuning were not significant.

Note that information about maternal education level was missing for n=5 TD children and n=7 children with ASD and exact gestational age at birth was missing for n=4 TD and n=4 children with ASD. None of the children for whom this information was missing were reported to have been born prematurely (<36 week gestation).

**Supplementary Figures**

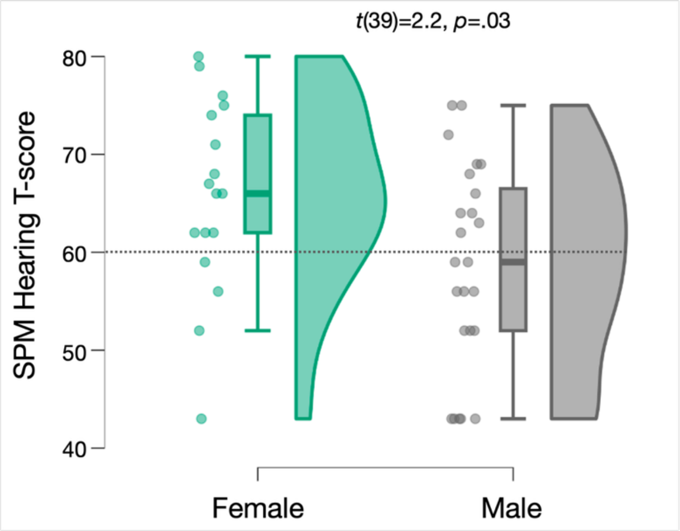


**Figure S1.** Sex differences in SPM Hearing T-score in children with ASD or other NDDs. Dashed line at a T-score of 60 indicates threshold for sensory processing difficulties.

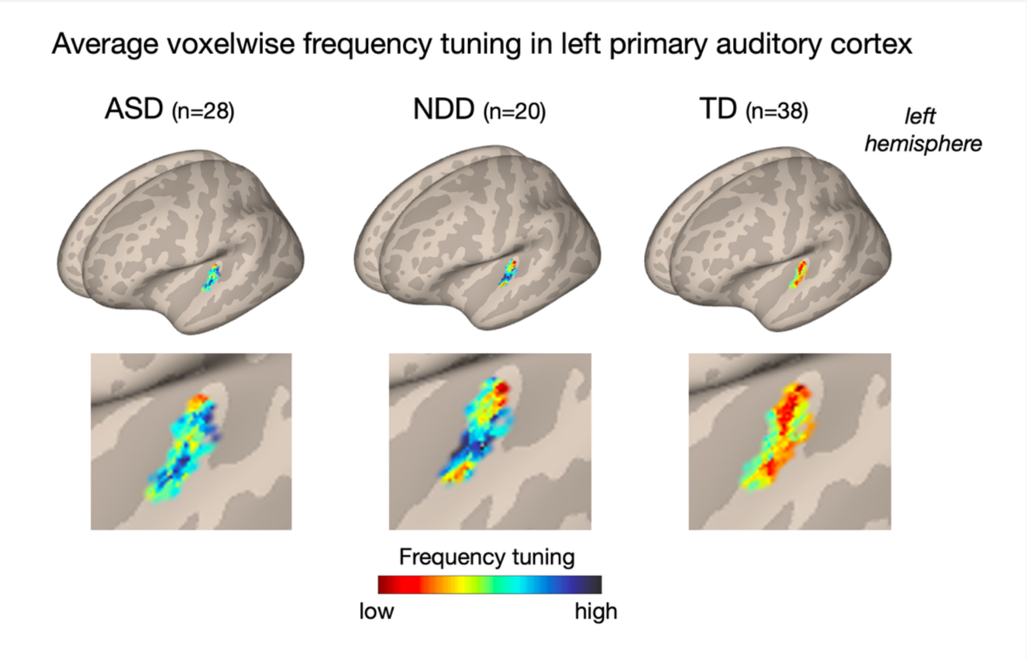


**Figure S2.** Maps of average frequency tuning for each voxel in left primary auditory cortex for each diagnostic group. Red colors reflect tuning to lower and blue colors to higher frequencies.
